# Zebrafish facility report on implementation of artificial plants as structural enrichment

**DOI:** 10.64898/2025.12.04.692061

**Authors:** Aymene Youcef Krachni, Richard Busch, Indigo Brakus, Angelina Schumann, Jenny Wilzopolski, Nils Ohnesorge

## Abstract

There is a broad consensus that husbandry conditions of laboratory animals need constant improvement to guarantee optimal animal welfare and research data quality. Zebrafish (*Danio rerio*) as one of the main animal models in biomedicine and toxicology are currently kept in barren tanks in most experimental setups, as well as in animal husbandry. Structural enrichment with artificial plants is currently discussed as a potential refinement measure to provide a more diverse environment. It was shown, that this can reduce stress and improve cognitive abilities, survival rate and fertility in these animals. Still, concerns remain regarding its long-term benefits and drawbacks. Therefore, we introduced artificial plants in our husbandry tanks, and evaluated over a one-year period if in our specific system, the benefits would outweigh the risks. Comparing six different lines of zebrafish raised in enriched tanks or under our standard conditions, we saw no significant difference in terms of sex ratio, fertility or pathogen burden. Enrichment increased the survival rate of 4 % during upbringing, albeit not statistically significant due to high variation. When analyzing zebrafish behavior in their 8 L home-tanks, we found less zebrafish in areas close to the plants but rather a preference for the open water in the middle or opposite side of the tank. This effect was even more pronounced at lower holding densities of less than 4 fish per liter. In summary, we found that introducing structural enrichment to our zebrafish facility carried low cost and no detrimental effects for the animals but at the same time benefits were difficult to determine in our readouts as our survival rates already were very high without these measures. We like to encourage other animal facilities to evaluate enrichment measures to ensure a broader discussion on long-term benefits of structural enrichment in zebrafish husbandry systems.

## Introduction

Zebrafish (*Danio rerio*) is among the most successful animal models worldwide. Publication numbers increased by 75% in the past ten years (according to PubMed with 2814 publications in 2014 and 4949 in 2024, search term “zebrafish”, accessed on 29^th^ September, 2025). Despite its popularity in many research areas like biomedicine and toxicology, few is known about the optimal husbandry conditions for laboratory zebrafish. In the past, mainly barren tanks were used to improve hygiene standards and efficacy. Nowadays, the importance of zebrafish husbandry conditions and especially the need for improvement is increasingly recognized, as reflected by a growing number of publications (Tsang and Gerlai 2024; Sen Sarma et al. 2023). The reasons are on the one hand ethical obligations for best possible animal welfare as formulated in the EU directive 2010/63. On the other hand the acknowledgement of the role of environmental factors on data quality and reproducibility (Paull, Lee, and Tyler 2024). Hence, researchers should aim for a better understanding of zebrafish needs. As a first step, European legislation was recently updated to include a restricted range of permitted water parameters, acknowledging this process (European Commission 2024).

A way for improving husbandry conditions could be by providing appropriate environmental enrichment with the aim to increase the animal’s environmental complexity via structural, dietary, social or cognitive measures (Näslund and Johnsson 2016). Structural enrichment is defined as modification of the animal’s habitat or the addition of physical structures or objects to promote natural behavior and increase overall well-being. Among others reduced anxiety and stress as indicated by lower cortisol levels, better memory, learning and survivorship were reported (Stevens, Reed, and Hawkins 2021; Buenhombre et al. 2021; Lee, Paull, and Tyler 2019). In contrast, effects on aggression differed among studies, where both an increase and a decrease were observed (Stevens, Reed, and Hawkins 2021). Despite these encouraging findings, structural enrichment is still lacking similar acceptance as other forms of enrichment like visual or social enrichment or use of live feed that are increasingly common (Lidster et al. 2017). An international survey among zebrafish facilities looking into this aspect reported widespread concerns regarding high workload, additional costs, consistency of scientific results and increased risk of pathogen growth. As a consequence, only the minority of zebrafish facilities have implemented structural enrichment or planned to do so (Lidster et al. 2017). Unfortunately, more recent data is lacking, but reported implementation in method sections of zebrafish publications remains low with a single mention of any form of refinement or enrichment in the 100 most recent research articles listed on PubMed employing zebrafish (search term “zebrafish”, accessed on 8^th^ July, 2025, see **Supplement 1** for details). Apparently, for many zebrafish facilities the feared risks or costs still outweigh the potential benefits and additionally the lack of reporting impedes change. One of the main problems is that the potential benefits are so far less tangible than improved water quality or better standardization of water parameters and that uncertainties remain if the findings of individual studies are transferable to standard husbandry settings. Some of these studies were conducted with single or pair-housed fish, that in addition have been removed from their home-tank and transferred to a novel environment (von Krogh et al. 2010; Keck et al. 2015), while others studies showed differences in enrichment preferences between single, paired or group-housed fish (Spence, Magurran, and Smith 2011; Schroeder et al. 2014). Therefore, findings on individual or paired-housed zebrafish might be more applicable to more stressful, experimental settings than for standard group-housed husbandry of a shoaling species.

So far, most studies have only investigated the effects of structural enrichment for a short period of time, e.g., a few days or weeks. The question of the long-term effects of such measures remains open. In the systematic review of Gallas-Lopez et al. on effects of structural enrichment only 3 out of 27 included studies were performed over a longer time frame than 3 months, while 21 were performed for less than a month (Gallas-Lopes et al. 2023). Therefore, it needs further investigation if structural enrichment maintains its positive effects over time or if constant changes are necessary to avoid habituation. Still, the conclusion of the systematic review was, that despite many confounding factors, the overall positive effects for structural enrichment are holding up (Gallas-Lopes et al. 2023).

Recent studies highlight that the form of the structural enrichment matters as well, as promising results were achieved with gravel substrate, artificial plants or similar simulations of vegetation while more artificial objects like an arrangement of glass rods or airstones were not beneficial or even counterproductive (Wilkes et al. 2012; Schroeder et al. 2014). In general, it has to be taken into account that structures inside the tank reduce free swimming space, might block the view on the fish for daily health inspection, could potentially inhibit water flow thus creating areas of low water quality or could induce territorial aggression between fish (Woodward, Winder, and Watt 2019). Therefore, the structural enrichment has to be provided in a way, that the benefits outweigh the risks (Lee 2022). Both real and artificial plants have been tested to improve the tank environment. As a general rule, artificial plants are preferable over live plants in regards of biosafety, handling and reproducibility but low-quality plastic material poses the risk of leaching chemicals to the water with hard to determine long-term consequences. In contrast, it was shown, that gravel pictures underneath the tanks were providing as good enrichment as real gravel inside the tanks, thereby avoiding most of the aforementioned problems (Schroeder et al. 2014). Observations from the natural habitat of the fish used should guide the decisions on how to structure the tank environment (Tsang and Gerlai 2024).

Overall, the challenge remains to find suitable readouts to determine the welfare state of the fish, beyond the absence of illness. Here, among others, methods to investigate choice preference, levels of stress or anxiety, improved learning and memory, exploration behavior, aggression or fecundity have been employed (Stevens, Reed, and Hawkins 2021; Lee 2022). As additional requirements for standard husbandry settings these methods should be non-invasive and performed in their home-tank to avoid any additional stress or burden for the fish. In this way, they can be carried out regularly as part of the welfare assessment. Survivorship and sex ratio, for example, are such parameters that are constantly recorded in animal facilities and are easily available to assess enrichment effects (Lee, Paull, and Tyler 2019).

In summary, zebrafish are a shoaling species, preferring open-water areas and that evolved with various forms of vegetation in their natural habitat. They are highly adaptable and can live under a wide range of conditions that helped to promote their status as one of the major research models. It is necessary to optimize their laboratory husbandry conditions and structural enrichment as a form of refinement that could improve animal welfare and research data quality. But widespread implementation is still lacking as published studies on its effects are still missing the robustness needed for long-term benefits to outweigh concerns for transferability to standard husbandry, pathogen risks and cost. To address these concerns and close the knowledge gaps, we documented implementation of artificial plants in our facility and compared its effects with our standard husbandry conditions in all forms of readouts that we have already established as considered relevant for our research. We documented the results of fish survival and sex ratio after rearing, mating success, pathogen burden in the water and behavior of fish in their home-tanks with or without structural enrichment over the period of one year. Therefore, our aim for this study was to investigate whether positive or negative effects of structural enrichment could be detected in our facility-specific setting under long-term husbandry conditions in home-tanks, without any additional burden for the fish and low additional cost or work-load for the facility.

## Methods

The facility report was prepared and conducted according to PREPARE, ARRIVE and FAIR guidelines (Smith et al. 2018; du Sert et al. 2020; Wilkinson et al. 2016).

### Animals

The following fish lines were used: *elavl3:H2B-GCaMP6s*^*jf5*^ (line 1a, 1b), Tübingen wildtype (WT TU, line 2a, 2b), *HuC:GCaMP5g*^*a4598*^ (line 3), *HuC:Gal4/UAS:RFP* (line 4), *mitfa*^*w2*^ (line 5) and Tüpfel-Longfin wildtype (WT TL, line 6). All lines except WT TL were maintained with Tübingen wildtype genetic background. In addition, *elavl3:H2B-GCaMP6s, HuC:GcaMP5g* and *HuC:Gal4/UAS:RFP* were in a *mitfa*^*-/-*^ background. No more than 3 generations of sibling matings were performed before lines were outcrossed against TU wildtype or *mitfa*^*-/*-^ lines with high genetic heterogeneity to avoid inbreeding depression. TU and TL wildtype stocks are newly re-imported from the European Zebrafish Resource Center (EZRC) every 3 years to avoid creation of sublines.

### Environmental parameters

Adult zebrafish were kept in 3.5 L or 8 L tanks with gravel pictures underneath, not segregated by sex at a holding density around 5 fish/L in a Standalone husbandry rack (Tecniplast, Active Blue) with copper-free, recirculating water and a daily water exchange of 10-20%. Reverse osmosis was used for tap water purification from copper free piping, followed by reconstitution with sea salt. Water parameters were maintained at 28 °C, pH 7.5, conductivity 800 µS/cm, oxygen levels >7 mg/L, carbon dioxide levels 1-6 mg/L, carbonate hardness 2-5 °dH, total hardness 2-6 °dH, ammonia <0.05 mg/L, nitrite <0.02 mg/L and nitrate <25 mg/L (Aleström et al. 2020). All water parameters were either monitored continuously (temperature, pH, conductivity), daily (oxygen, carbon dioxide) or at least once a week (carbonate hardness, total hardness, ammonia, nitrite, nitrate). Room temperature was 24-25 °C, light intensity 500 lux in front of the tanks and 10-20 lux at the back of the tanks. Illumination was set to 10 hours of darkness followed by 30 min of dusk with increasing intensity to the maximum of 500 lux for 13 hours, followed by 30 minutes of decreasing light to darkness again. The adult fish were fed twice daily dry food (Sparos 400-600) *ad libitum* and once per day artemia (Sanders). Health checks were performed daily during feeding.

### Structural enrichment

Pilot experiments were performed to determine the most promising form and position of structural enrichment. Different plant forms and sizes were acquired that were certified to be made of non-leaching polyethylene. Four of these plants were selected that are present in the natural habitat of zebrafish and which were not too large or spiky to fit in our tanks as well as not to pose a danger of injury or swimming impediment to the fish (Dohse Aquaristik, *Hygrophila* #51559, *Vallinesria* #41507, *Rotala* #41514, *Bacopa* #51555). The base of the plants was manually removed as the connecting glue to attach the plants could not withstand hot water for long during the cleaning procedures and might contain leaching chemicals. Instead, a random mix of plastic plants weighing in total about 5 g and 5-10 cm length were attached to the left tank wall with a rubber suction cup. They were positioned in a way that they stand upright in the water column, that water flow through the tank was not inhibited and no food remains would gather around the plants (**Figure 1**). In general, the tanks remained easily accessible for cleaning or netting fish. Either one plant bundle was attached in a 3.5 L tank or two next to each other in an 8 L tank. Every 1-2 months, when a tank needed more thorough cleaning and biofilm removal, fish were transferred to a new tank and received a new plant bundle of similar size but slightly different composition to avoid habituation.

**Figure 1:**
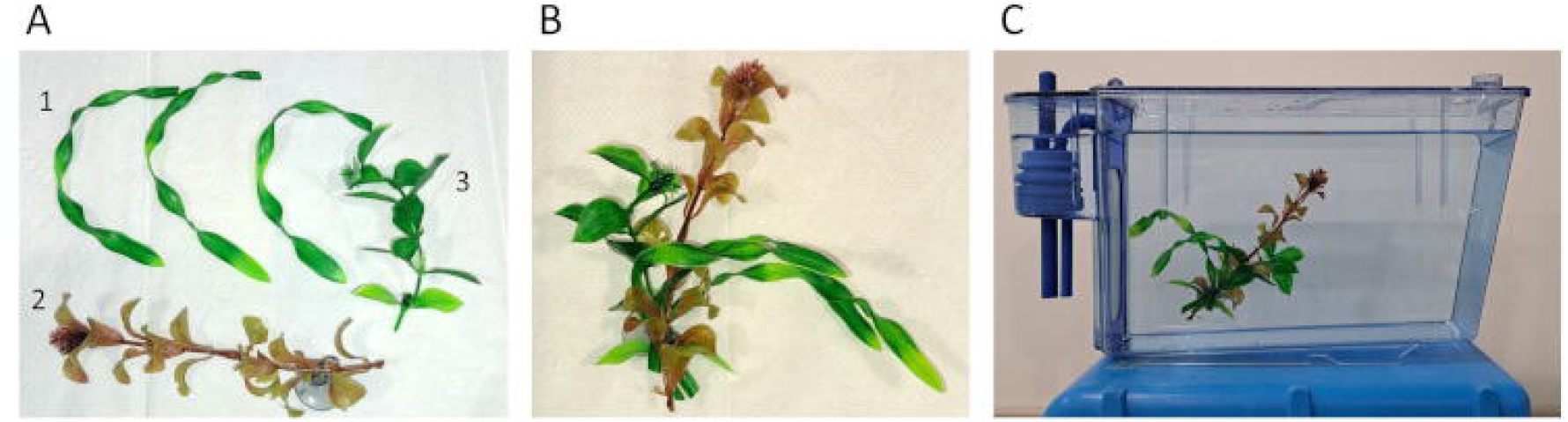
Artificial plants as structural enrichment. (A) A combination of artificial plant parts recreating in this example *Vallisneria* (1), *Rotala* (2) and *Hygrophila* (3) were bundled (B) and attached to the tank wall using a rubber suction cup (C).

### Rearing

All fish lines used were generated in our quarantine system and needed bleaching of the eggs before being transferred to the main system for rearing. In brief, bleaching was performed two-times for 5 minutes with 60 ppm sodium hypochlorite based on the ZIRC protocol of “Embryo Surface Sanitation” with E3 medium for washing instead of E2 medium. Larvae were checked under a stereo microscope (Olympus SZX16/SZX2-ILLT) to be inconspicuous in morphology and behavior and were transferred from petri dishes to the main system at 5 days post fertilization (dpf). If sufficient larvae were available for rearing either 25 larvae were raised in a 3.5 L tank or 50 larvae in an 8 L tank. All tanks had gravel pictures underneath. Larvae were initially fed only with dry feed (Sparos 100-200), then with increasing fish size feed of larger particle size was added or replaced the smaller sized feed (Sparos 200-400, 400-600). Dropwise artemia feeding started at 10 dpf with increasing amounts during rearing. All lines were raised in the same husbandry rack over the course of a year. They were evaluated for survival, sex ratio and their first two mating results. Tanks of the same line with or without structural enrichment were placed next to each other on the same level in the husbandry rack. The survival rate and sex ratio were determined when the fish were first used for mating at the age of 3 to 6 months.

### Mating

Fish aged 3 to 6 months were used to study effects of structural enrichment in home-tanks during rearing on the first two matings. For mating 2 females and 1 male were randomly selected and placed into a 1 L mating tank at 4-5 PM with both sexes separated. The mating tanks were then placed on gravel pictures overnight. On the next day at 8.30-9 AM tank water was exchanged and dividers removed. The tanks were placed skewed on gravel pictures to create areas of shallow and deep water to simulate the shore and facilitate haptic stimulus. Mating was allowed to continue for one hour. After the mating the fish were returned to their home-tanks. The eggs were collected in a strainer and then counted and distributed to 1x E3 embryo medium containing petri dishes with up to 50 eggs per 10 cm dish in 20 mL E3 medium. 1x E3 medium contains 5.14 mM NaCl, 0.18 mM KCl, 0.34 mM CaCl_2_ and 0.41 mM MgCl_2_ and was prepared in 60x stock solution with a pH titrated to 7.2. Larvae were kept for up to 5 days in an incubator set to 28.5°C with 14/10 light-dark hours.

To investigate effects of structural enrichment added to mating tanks only Tübingen wildtype fish were used, aged 4 to 9 months. On each occasion three tanks were set up for mating as described above, but separated with laminated cardboard between the tanks to prevent visual interaction between fish in neighboring groups (**Figure 2**). From these three tanks one tank was placed on an isochromatic colored beige-sand colored tray, the second tank was placed on a gravel picture and the third tank was both placed on a gravel picture and a small, swimming plant part was added for enrichment. After 1 hour of mating the fish were returned to their home-tanks and eggs were counted to determine the clutch size.

**Figure 2:**
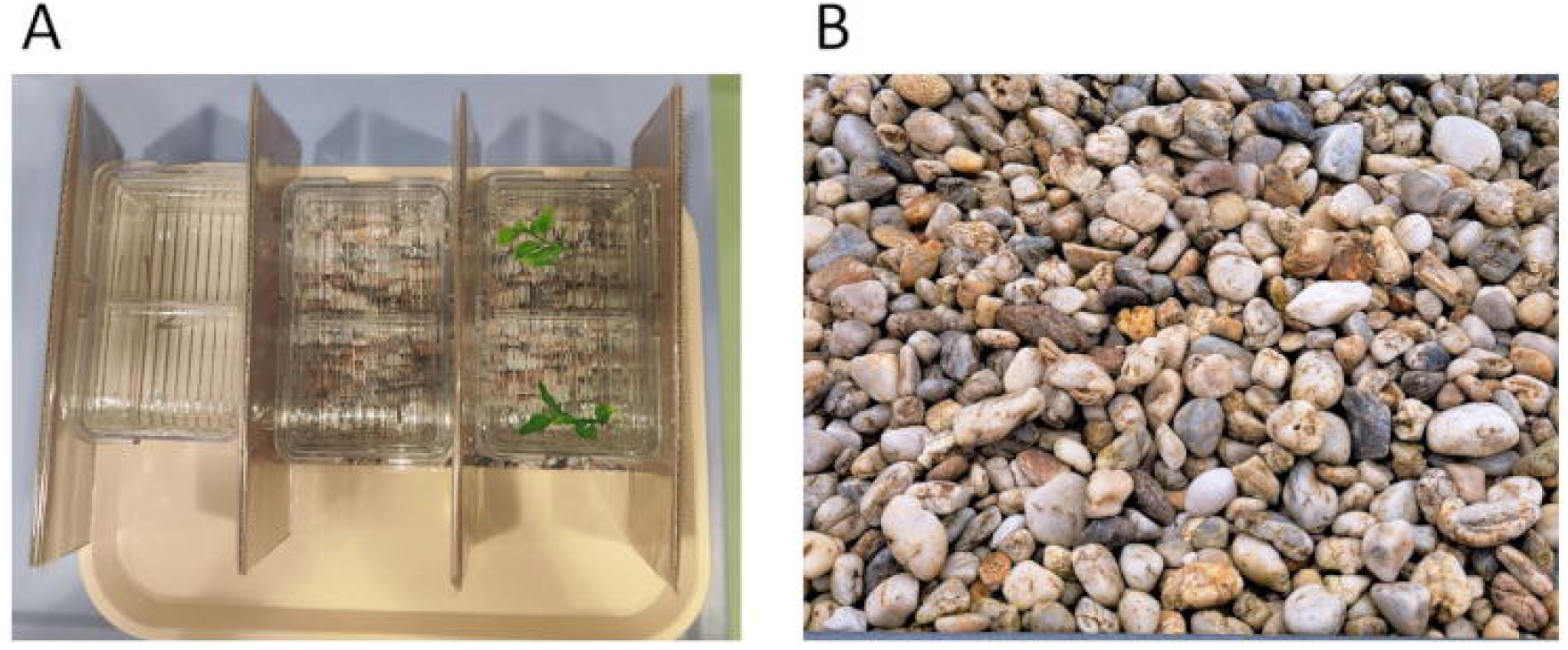
Structural enrichment during mating. (A) The effects on mating success and numbers of eggs laid were evaluated using three different conditions for mating tanks that were separated by laminated cardboards: tanks without enrichment, gravel picture only or gravel picture and small plant (from left to right). (B) Shown is the gravel picture used for structural enrichment during mating, which was printed in DINA4 and laminated.

### Observation of fish/plant interaction

To determine the effects of the structural enrichment on the behavior of the fish, their place preference in the tank with or without artificial plants was investigated. In addition, two situations were distinguished with either ongoing human activity in the husbandry room or with undisturbed fish. Both situations take up roughly 50% of the 13 hours daylight time in our facility on average with more activity in the morning to early afternoon and the undisturbed phase in the late afternoon and evening.

To this end, images of 8 L tanks with fish aged 4 to 13 months were taken from a 1.5 m distance in front of the racks. In the first case with human presence representing ongoing activity in the facility, pictures were taken directly at 4 PM. To observe behavior of undisturbed fish a camera was set up at 4 PM and images were taken automatically with one frame per minute starting 1 hour after the last person left the facility from 5 PM to 9.30 PM. In both cases the positions of fish were then identified and highlighted as region of interest in these pictures and they were digitally overlaid with a template to separate the tank equally either in left and right or in left, middle and right areas. Number of fish per area were counted. In case a fish was located on the border between two compartments, the compartment was chosen where the larger part of the fish body was located. In case equal parts of the fish were in both sections, the one where the head was located was chosen. From an initial pilot experiment an effect size of 10% was estimated and under the conditions for a power of 90% and an accepted alpha error of 0.05 it was calculated that a sample size of 34 images was needed. With 4 tanks per condition available, either individual pictures were taken on 9 different days for final evaluation of directly observed fish, or 9 pictures were extracted from the movie of undisturbed fish with each 30 minutes apart. In general, all fish swam calmly and steadily and showed no obvious signs of stress or disturbances when the pictures or movies were taken. If not mentioned otherwise, 26 to 31 fish were present per 8 L tank resulting in a holding density of 3.3 to 3.9 fish per liter.

Different survival rates of lines 3, 4, 5 and 6 resulted in different holding densities for these lines. This allowed for investigation of its effects on place preference. Fish of these lines were initially separated during upbringing in two 3.5 L tanks with and without enrichment. At the age of 3-4 months, they were used for matings and then united in a single 8 L tank with enrichment. 1-3 months later pictures and movies were taken to analyze their place preference.

### Total microbial count

Total microbial counts are performed in our facility on a regular basis to investigate the hygiene status via water samples from the sump. Additional samples were taken from the back left corner of lines 1a and 1b holding tanks once per week on three occasions to check for an increased risk of pathogens due to structural enrichment. The samples were diluted by 1:10 and 1:100 with sterile ddH_2_O to achieve a countable range of <300 colonies. Sterile water was used as control. 0.1 mL of each diluted sample and control was plated to 20 mL sterile PCA (plate count agar) plates (20.5 g/L DM 195, Mast Group #121950) and rested upright for 15 minutes at room temperature until the liquid was absorbed by the plate. The plates were then incubated inverted for 72 +/-3 hours at 28°C, colonies were counted and number of colony forming units (CFU) per mL were calculated. Experiments with values for water control above 25 CFU/mL were rejected and repeated. Values below 10.000 CFU/mL were considered as safe and below 100.000 CFU/mL as acceptable in regards of fish health and welfare (Leonard, Blancheton, and Guiraud 2000).

### Statistics and data analysis

Power calculations were performed based on pilot data to estimate sample size number needed with the NC3R online tool (https://eda.nc3rs.org.uk/experimental-design-group) that used R 3.5.2 and the package power.t.test. In all cases an accepted alpha error of 0.05 and power of 0.90 was selected.

GraphPad Prism 10.1.2 software was used for statistical analysis. Gaussian distribution was tested via Shapiro-Wilk normality test and visually confirmed in a QQ plot. No outliers were identified in our data sets, tested by ROUT method with a maximum desired false discovery rate of 1%. Statistical significance of survival rates, sex ratio and mating success were calculated in a paired t-test. Significant differences between egg numbers were either calculated in an unpaired t-test for the same mating conditions after enriched upbringing or with a One-way ANOVA to test for different enrichments during mating. Differences in place preference between left and right tank side were determined in an unpaired t-test. Statistical significance between place preference of left, middle or right area was calculated via Two-way ANOVA followed by multiple unpaired t-tests or Tukey’s multiple comparisons test.

### Ethical approval

As part of our 3R strategy for reducing surplus animals only fish lines already required for other projects or line maintenance were used to study effects of structural enrichment. In addition, only readouts that did not put additional burden beyond necessary husbandry practices on the fish were selected. Refinement experiments were exempt from approval by local German authorities (LAGeSo) but the zebrafish husbandry was authorized (ZH 181). Zebrafish were kept in accord with EU directive 2010/63/EU on the protection of animals used for scientific purposes.

## Results

Over the course of one year six different lines have been raised in 16 tanks in the same husbandry system. Of every line half the tanks received structural enrichment in the form of artificial plants while the other half were kept under our standard conditions. A detailed results table with fish numbers and lines is given in the supplements (**Supplement 2**).

Historically the survival rate for zebrafish in our facility determined after rearing at 90 dpf has been 73 % on average in the two years before the implementation of structural enrichment started. This value was closely matched with an average survival rate of 74 % for fish reared again under these same standard conditions analyzed in this study. On 6 out of 8 occasions the fish under enriched conditions had a better survival rate compared to their siblings. On average the survival rate was about 4 % better without being statistically significant (**Figure 3 A, B**).

**Figure 3:**
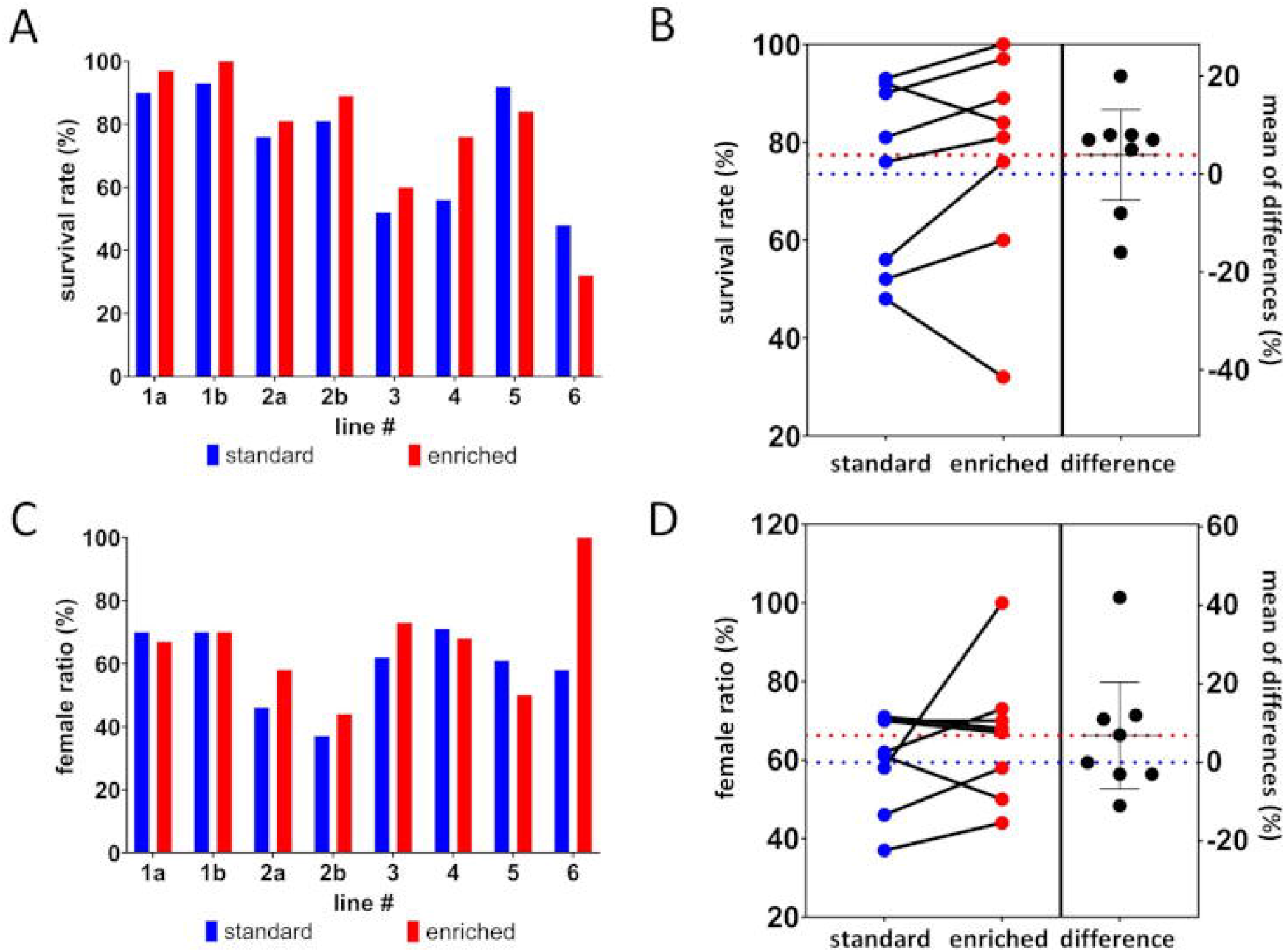
Influence of structural enrichment on survival rate and sex ratio during rearing. Sixteen tanks of 6 different lines were evaluated around 90 dpf after rearing with or without structural enrichment. (A) In 6 of 8 cases the survival rate was better with structural enrichment than the control tank, albeit on average not statistically significant (p = 0.353). (B) The average survival rate for standard conditions was 74 % (blue dotted line) and for enriched conditions 77 % (red dotted line). A pairwise comparison of the individual tanks (enriched minus standard) showed differences in survival of -16 % to +20 %, with an average difference of 4 % (dotted lines). (C) Ratio of females per tank and line were determined at the first two matings. (D) No significant difference could be determined regarding the sex ratio with an average of 60 % and 66 % of females per line for standard or enriched rearing respectively (dotted lines). The pairwise comparison of enriched tanks and their control counterparts showed no significant change as well.

The sex of the zebrafish is determined mostly by environmental factors during upbringing and structural enrichment might reduce the available swimming space and thereby the perceived holding density for the fish. The sex ratio was determined at 90 dpf or later during the first two set ups for matings. In case of line 6, only females developed under enriched conditions. This was probably due to the low survival rate resulting in a low holding density which is known to favor female sex determination (Ribas et al. 2017). Overall, no significant difference was found with an average of 60 % and 66 % females per line raised under standard or enriched conditions respectively (**Figure 3 C, D**).

One reported benefit of structural enrichment is the reduction of both stress and anxiety for the fish. We hypothesized that this could result in increased numbers of successful matings and egg production, especially on the first two occasions when the fish are not used to the handling and new environment of mating tanks which could potentially increase their stress response. Therefore, fish from the same line that were raised under standard or enriched conditions were set up in parallel when needed for mating. On average the zebrafish raised under enriched conditions showed no better performance in their first two matings with 61 +/-24 % and 82 +/-19 % of tanks having successful matings compared to the 64 +/-20 % and 75 +/-20 % of tanks with fish raised under standard conditions (**Figure 4 A, C**). Similarly, no significant difference was seen in the number of eggs laid on these occasions, with on average 68 +/-55 and 122 +/-58 laid from enriched fish compared to 72 +/-43 and 127 +/-56 eggs laid from not enriched zebrafish (**Figure 4 B, D**).

**Figure 4:**
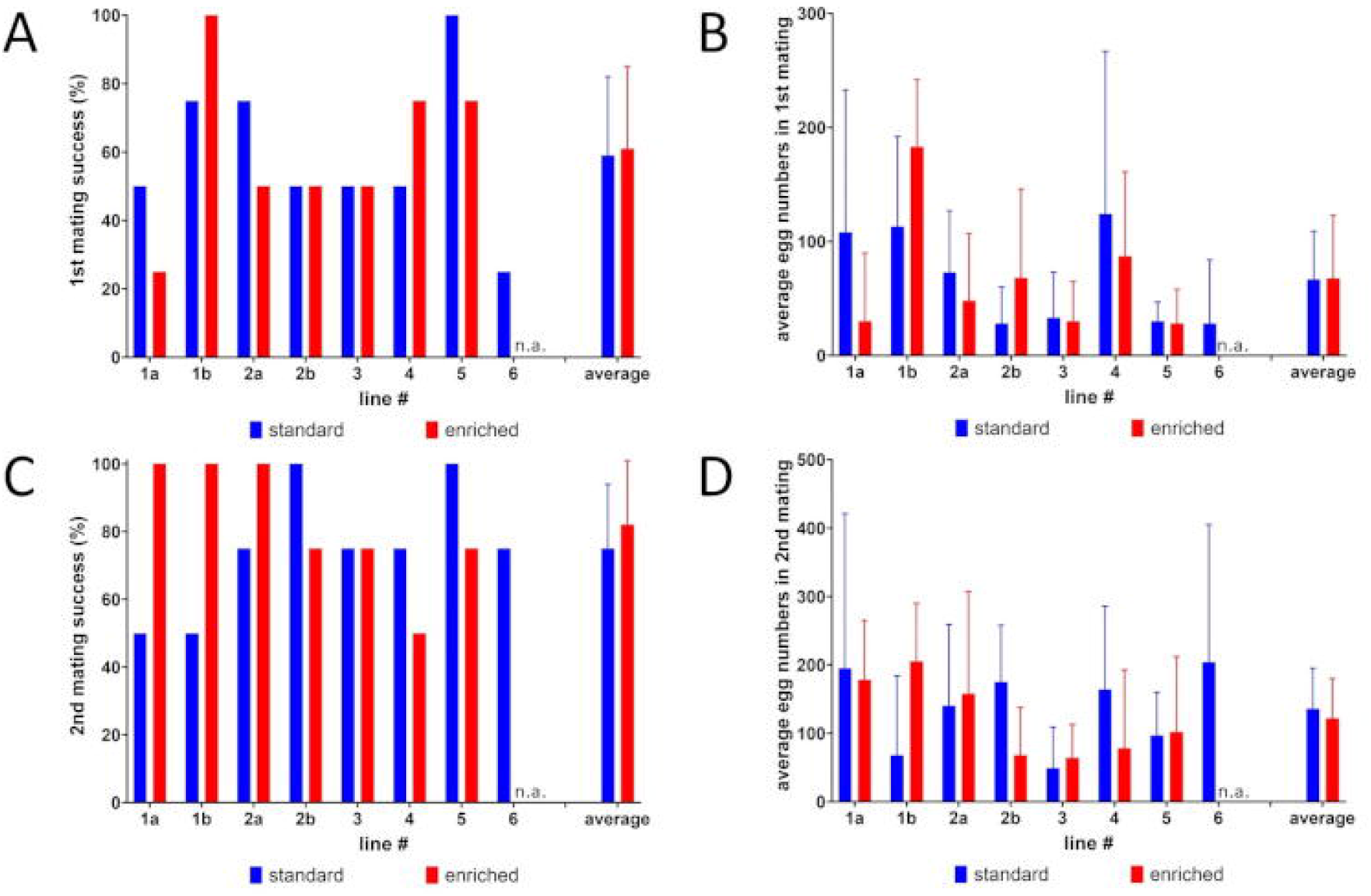
Influence of structural enrichment on mating success and egg production. After the fish reached adulthood, the first two matings that were performed on these lines were evaluated based on differences between standard (blue) or enriched (red) conditions during rearing. Only female fish were identified for enriched line 6. Therefore, no results were available for mating success and egg numbers (n.a.). Both in the first mating (A, B) and in the second mating (C, D) no significant differences were seen regarding mating success (A, C) or average number of eggs produced per mating (B, D). For the average values the standard deviation is indicated with error bars, n = 7-8.

Next, we evaluated if zebrafish in enriched tanks actually showed a difference in their place preference behavior compared to the non-enriched condition. For this, two scenarios were distinguished: About half of the daytime there is ongoing human activity in our facility, with standard husbandry procedures, fish matings and preparation for experiments. In general, the fish don’t show any forms of stress in regard to ongoing facility work but tend to stay closer to the front of tanks out of curiosity or in expectation of feeding. When lines 1 and 2 were analyzed for their place preference under these conditions, they evenly occupied both the left and right side of the tank under standard conditions (49:51) but avoided the left side with an artificial plant present (36:64) (**Figure 5 A-C**). On closer inspection, zebrafish preferred the open water of the middle area in a tank over wall sides under standard conditions (24:48:28) but rather moved closer to the right wall than to the plant under enriched conditions (14:45:41) (**Figure 5 D**). Similar results were seen for undisturbed fish, less pronounced but still significant with a left:right ratio of 44:56 and 20:47:33 for left:middle:right under enriched conditions (**Figure 5 E, F**).

**Figure 5:**
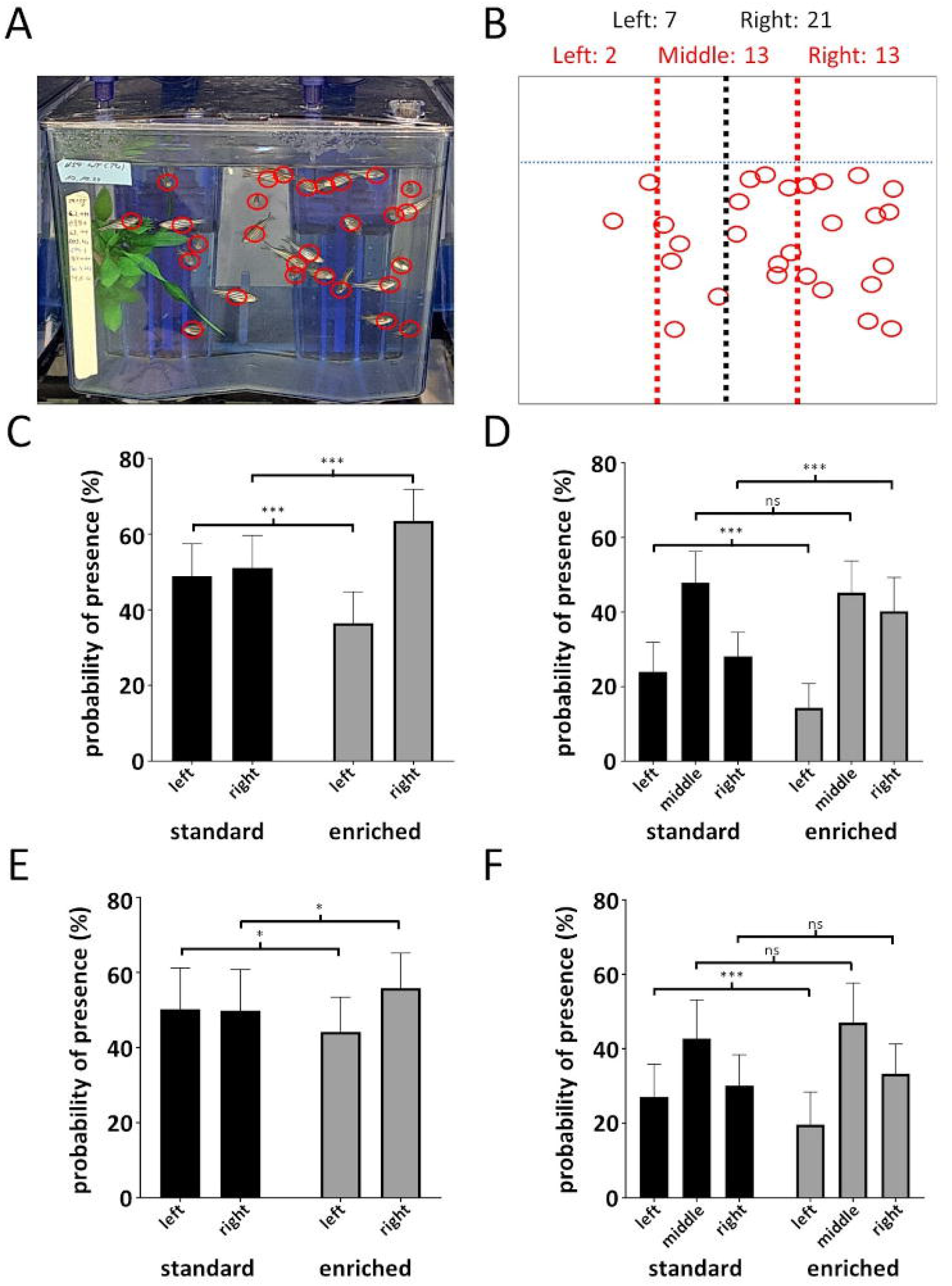
Effects of structural enrichment place preference. Lines 1 and 2 were investigated for their place preference. (A) One representative image is shown of an enriched home-tank, taken to identify the positions of fish. (B) Fish positions (red circles) were overlaid with a digital template to equally divide the tank either in left and right (black dotted line) or in left, middle and right (red dotted lines) areas. The dotted blue line indicates the water level. Then fish per area were counted. (C) With human presence and standard conditions fish are divided 49:51 between left and right side, while when enriched, they prefer the free area over the plant 36:64. (D) Divided in three areas the fish prefer the open water in the middle over the left or right area with walls (24:48:28) under standard conditions. Under enriched conditions they avoid the left area with the plant and stay more closely to the right wall (14:45:41). (E) When undisturbed by human presence, fish are divided 50:50 between left and right side under standard conditions, while with a plant present, they again prefer the free area over the plant (44:56). (F) Of the three areas the undisturbed fish under standard conditions prefer the open water in the middle over the left or right area with walls (27:43:30). Under enriched conditions they avoid the left area with the plant, too (20:47:33). Error bars indicate standard deviation, n = 36, ns = not significant, * p < 0.05, *** p < 0.001.

Not all fish known to be in the tank could be identified in a single picture but the amount of fish detected and counted was unchanged due to the presence of artificial plants with 83 +/-9 % for both standard and enriched tanks when directly observed (**Figure 5 C, D**) or 74 +/-12 % and 75 +/-9 % when video recorded under standard or enriched conditions respectively (**Figure 5 E, F**).

Different survival rates of lines 3, 4, 5 and 6 resulted in different holding densities of 3.5, 4.1, 5.5 and 2.5 fish/L, respectively. Analysis of place preference in enriched tanks under direct observation showed that in tanks with fewer fish and lower holding density the relative avoidance of the left side with the artificial plant was even stronger than for fish at higher holding densities (**Figure 6 A)**. Only 14% and 7% of fish at densities of 2.5 or 3.5 fish/L stayed in the plant area while at densities of 4.1 or 5.5 fish/L it was 28% and 23%, respectively. In contrast to what was seen before, a majority of over 50% of fish at low holding densities rather stayed close to the right wall, while at higher densities it was only 30-35%. This was also seen with the same fish when they were undisturbed and videotaped, although again to a lesser extent (**Figure 6 B**). In this case, only the fish with the lowest stocking density of 2.5 fish/L showed a different behavior than the others, as with 43% a significantly larger part of them stayed near the right wall, compared to 31-34% of the other lines.

**Figure 6:**
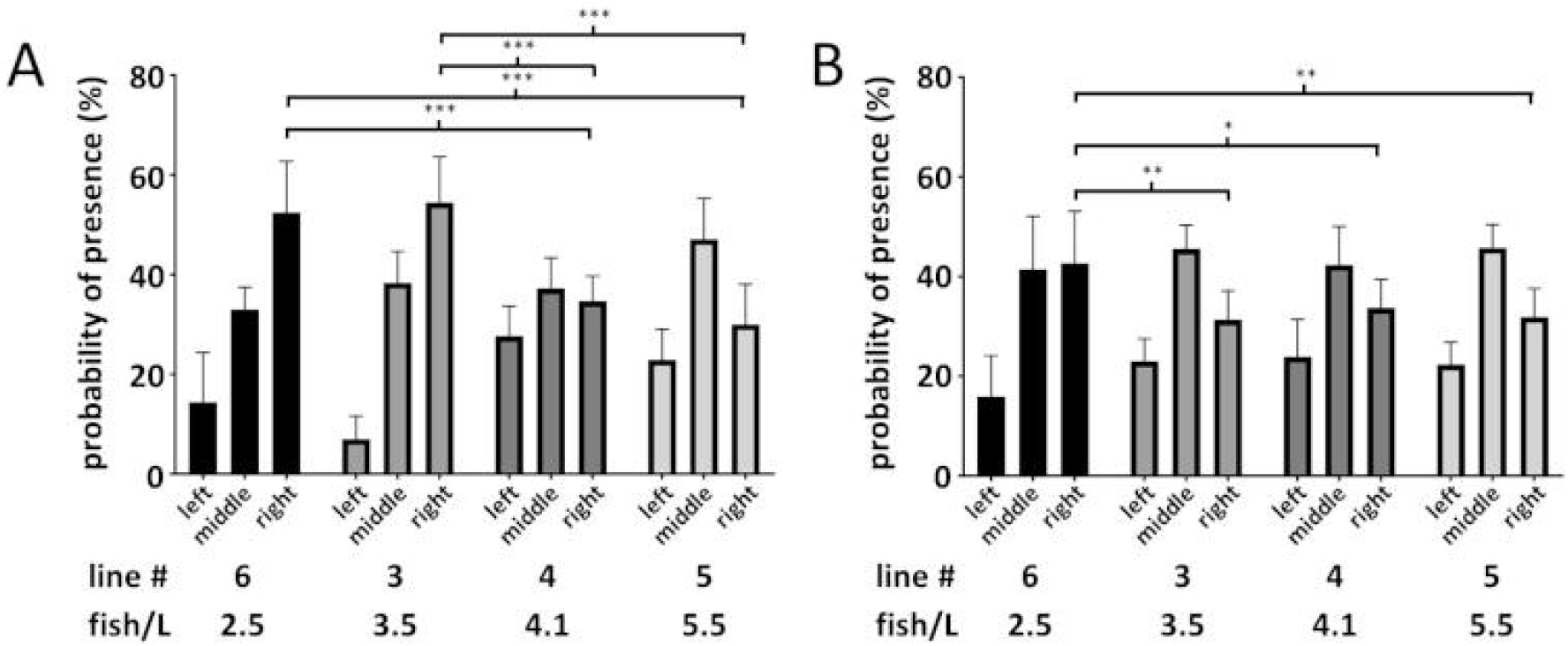
Effects of holding density on place preference. Enriched tanks of lines 3, 4, 5 and 6 with different holding densities (fish/L) were observed for place preference. (A) With human presence fish with lower holding densities of 2.5 or 3.5 fish/L had an increased avoidance of the enriched left tank side and a stronger preference for the right wall side. (B) Undisturbed fish with the lowest holding density of 2.5 fish/L still preferred more the right wall side compared to tanks at higher holding densities. Error bars indicate standard deviation, n=9, * p < 0.05, ** p < 0.01, *** p < 0.001.

Another concern with structural enrichment is the increased risk for pathogens. After two months of continuous use, we did not observe biofilm growth on the artificial plants but instead on the comparatively small area of the suction cup. In addition, we placed them in a way that they most likely did not interfere with water flow, as remains of feed and feces could quickly facilitate exponential bacterial growth. To assess this risk, we analyzed water samples from holding tanks of line 1a and 1b from three different occasions in regard to their microbial burden via a total microbial count (**Figure 7 A**). The mean number of colony forming units (CFU) per mL were 4633 and 5673 for line 1a and 2387 and 14620 for line 1b with or without enrichment, respectively (**Figure 7 B**). In a repeated measures One-way ANOVA no significant difference was found (p = 0.446). Overall, the values were in the same range as our whole holding system, with a mean of 8843 CFU/mL in the last three measurements of samples from the sump.

**Figure 7:**
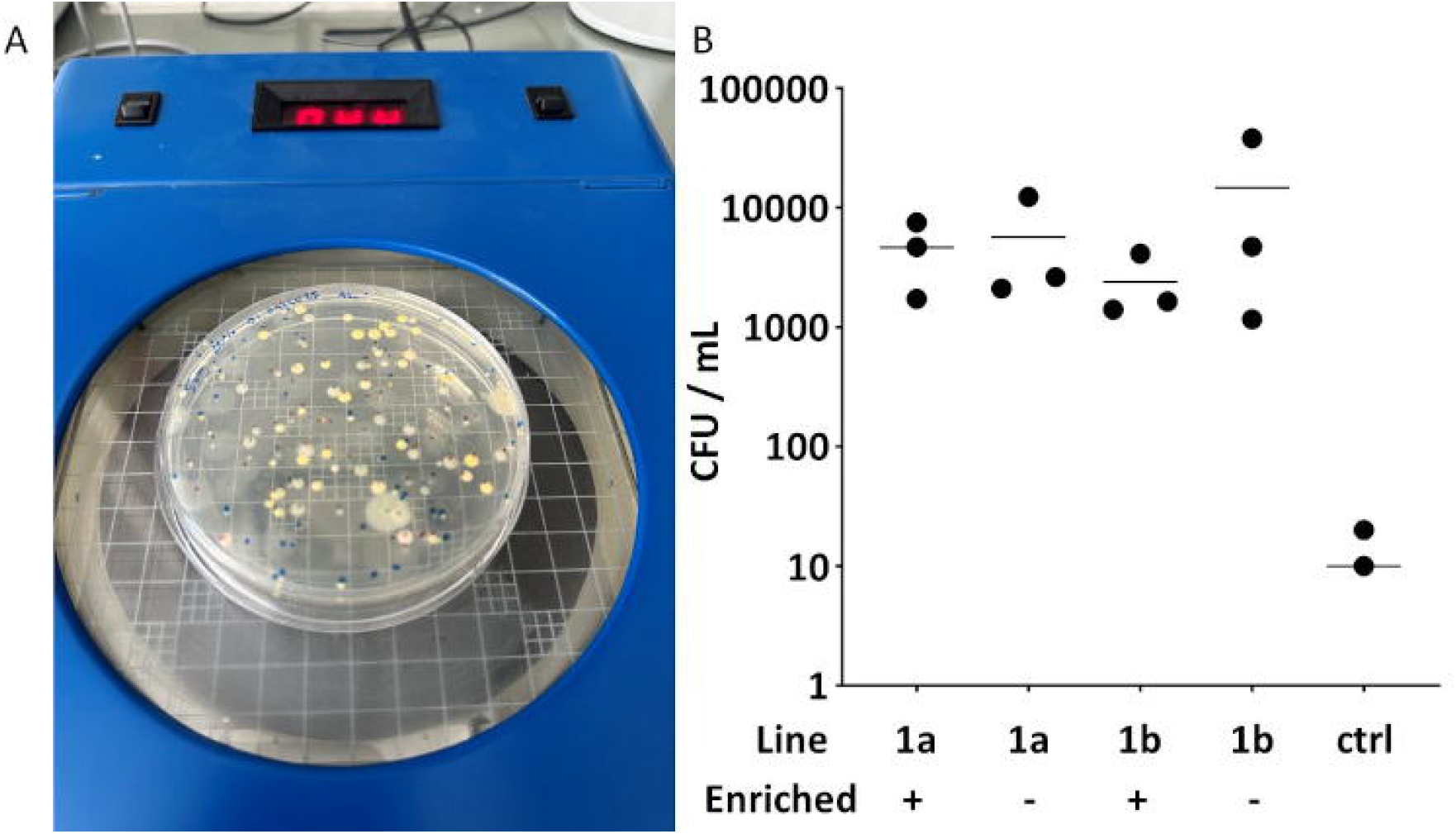
Effects of structural enrichment on the risk of pathogens. (A) Water samples were taken from the left bottom corner at the back of the tanks of lines 1a and line 1b on three different occasions and plated on PCA to determine the total microbial count. Sterile water was used as control. (B) No significant difference in CFU/mL was observed in tanks with (+) or without (-) enrichment. The third count for the water control (ctrl) was 0 which cannot be displayed on a logarithmic scale. The horizontal bars indicate mean values, n=2.

Lastly, we wanted to investigate the effect of structural enrichment during mating as another regularly occurring husbandry procedure. In contrast to the former experiment (**Figure 4**), now all fish were taken from the same home-tank irrespective if there was enrichment or not and randomly distributed to three different tanks with standard conditions, gravel picture or gravel picture plus plant (**Figure 2**). Only fish of line 2a and 2b (Tübingen wildtype) were used. Here, on average 149, 124 or 135 eggs were laid per mating for standard conditions with an isochromatic beige-sand underground, enrichment by gravel picture under the mating tank or an artificial plant inside the tank in addition to gravel pictures, respectively (**Figure 8**). No significant differences were found based on calculation of repeated measures One-way ANOVA between any of these conditions (p = 0.564).

**Figure 8:**
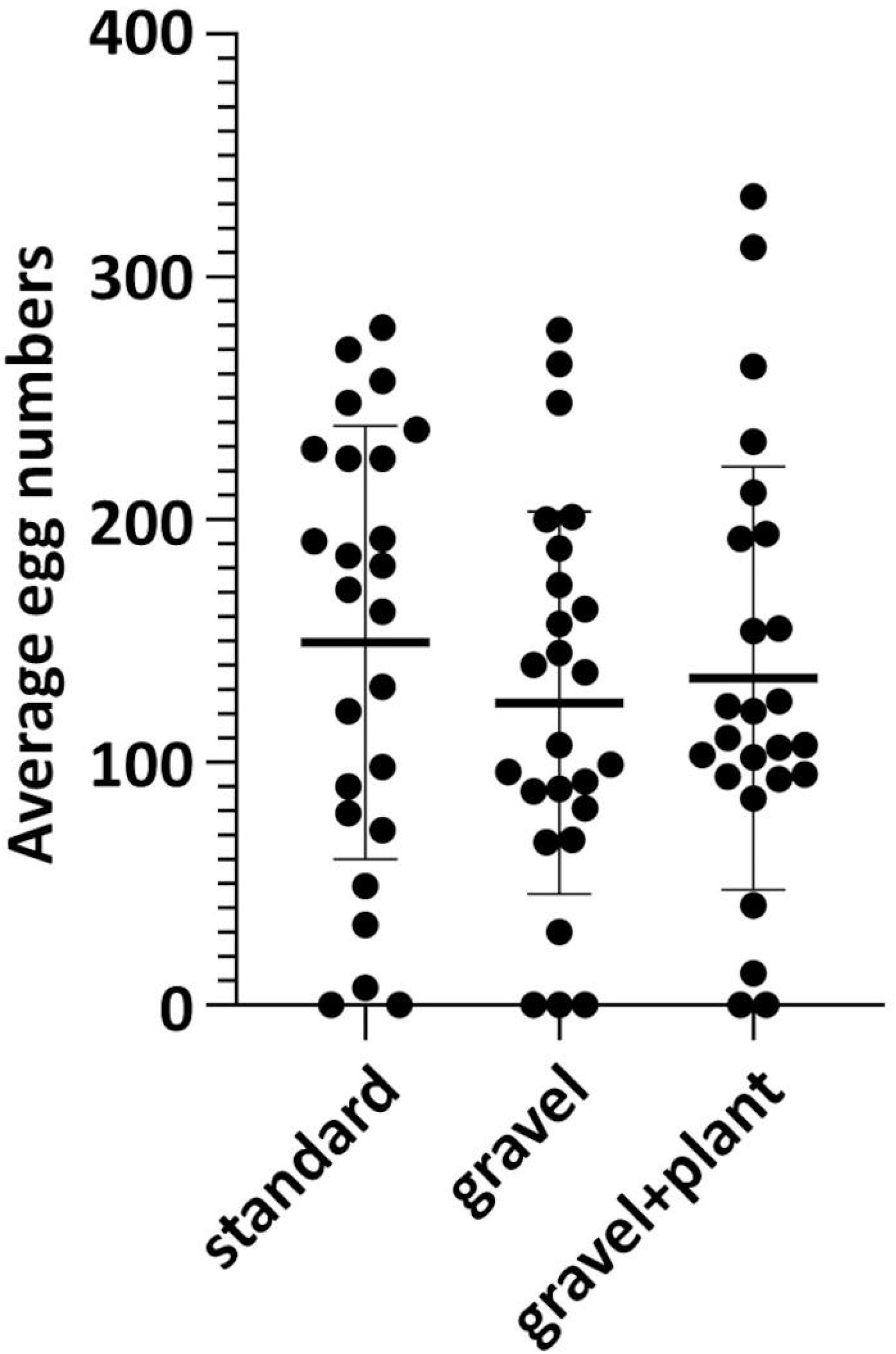
Influence of structural enrichment during mating. Wildtype Tübingen fish were repeatedly set up for mating with different enrichments and the resulting number of eggs were counted. On average neither the gravel picture under the tanks alone or together with artificial plants in the mating tanks resulted in a significant change of egg numbers received during matings. Error bars indicate standard deviation, n = 25 per group.

## Discussion

Aside from ethical and legal obligations there is consensus in the scientific community that laboratory animals need constant improvement of husbandry conditions to allow for optimal data quality (de Abreu, Parker, and Kalueff 2024). In addition, lack of standardization is also a concern for the reproducibility of research data generated with fish under different environmental conditions (de Abreu, Parker, and Kalueff 2024). It has been argued that structural enrichment actually increases data robustness because fish are more resilient even when less standardized objects are used (Tsang and Gerlai 2024). In the case of zebrafish, there is an ongoing discussion if artificial plants in husbandry systems could improve animal welfare. Currently only a minority of husbandry facilities use structural enrichment or are at least not reporting it according to the latest survey of Lidster et al. and the 100 most recent zebrafish publications we reviewed (Lidster et al. 2017). Named reasons were concerns regarding unclear benefits, costs, risks of pathogens, chemical leaching or increased aggression (Lidster et al. 2017; Woodward, Winder, and Watt 2019). For structural enrichment to be widely accepted and implemented in husbandry facilities, the benefits need to clearly outweigh the drawbacks. Nevertheless, the improvement of fish welfare can be challenging to measure directly. In addition, long-term effects are more relevant for husbandry but currently only few studies addressed these (Gallas-Lopes et al. 2023). Still, the first systematic review on effects of structural enrichment for zebrafish found that despite many confounding factors, overall, the positive effects are holding up (Gallas-Lopes et al. 2023).

Therefore, we decided to implement artificial plants as a new form of structural enrichment in our zebrafish facility and to monitor its effects in comparison to our standard husbandry setup for one year. We have chosen plants made of certified non-leaching polyethylene, with low material costs of 3-5 Euro per tank, adding up to less than 100 Euro for the whole rack, depending on the mix of plants used. After one year of more or less continuous use and including several dish washer cleanings at 80 °C, the material showed no sign of deterioration and we expected the plants used to last for several years. We were able to place the plants in the tanks in a way that they didn’t obstruct water flow or daily inspections of the fish, as demonstrated by similar detection rates for fish in pictures and movies taken from the tanks during place preference analysis. We found no increased biofilm growth on the plants and couldn’t detect an increase of the total microbial count in the water of the enriched tanks, indicating that UV light disinfection of our holding system was sufficient to keep microbial burden low (Rurangwa and Verdegem 2015; Leonard, Blancheton, and Guiraud 2000). Overall, we concluded that the chosen form of enrichment was low cost, easy to implement and not acting detrimentally on the water quality of our system or to the health of the fish.

We have not systematically investigated the effects of territorial behavior or aggression on the fish. We have occasionally observed that single fish remained near the plants and chased other incoming fish away, indicating a high place preference for these individual fish. This behavior never lasted until the next day and in the meantime didn’t seem to affect the behavior of the remaining shoal that kept calmly swimming in the open water area. There have been conflicting reports on how structural enrichment affects aggressive behavior and it needs to be further investigated in the future what combination of factors like holding density or the exact form of enrichment are inducing or preventing aggression (Woodward, Winder, and Watt 2019; Bhat, Greulich, and Martins 2015; Basquill and Grant 1998; Carfagnini et al. 2009; Sen Sarma et al. 2023).

From our experience, the main drawback for implementing structural enrichment was a small but unavoidable additional work load for our animal caretakers which we estimated for a complete rack with 50 enriched tanks to be one person-hour per month. This includes preparation or replacing appropriate structural enrichment, its maintenance in the tank, like re-attaching loosened structures or rearranging structures inside the tank during cleaning and cleaning the plants after use.

In summary, we found no significant detrimental effects by implementing structural enrichment in our facility, thereby creating a very low bar to be outweighed for potential positive effects for improved animal welfare.

For these, we used mainly readouts that are most important for our work as a facility and were therefore already established and continuously evaluated anyway. When comparing our new lines raised in regards of survival, sex ratio, mating success and numbers of eggs laid with or without enrichment no statistically significant effect was found.

In terms of survival, we saw a trend to better survival rates with enrichment as in 6 out of 8 cases survival rates were improved. The survival rate was especially low in the last case observed (line 6). This was likely due to a bad quality of this clutch as the parent line was already old and had several unsuccessful matings before a small number of eggs was obtained. These eggs were below standard quality and had to be raised nonetheless to maintain the line. This case might not be comparable with the other lines but we decided against excluding its data as it represents usual problems in a husbandry facility. Unfortunately, this resulted in a standard deviation to be much higher than initially anticipated and therefore we were unable to gather enough samples to reach statistical significance. This is in contrast to a considerable improvement of 29 % in the survival rate reported by Lee et al. 2019, which was likely only possible as only a single clutch was investigated. It has to be considered that in their study the survival rate of their standard conditions was only 54 % and with enrichment it reached 83 %, which is a similar level we have seen in our lab with standard conditions (Lee, Paull, and Tyler 2019). In general, as we already had established very high hygiene standards, excellent water quality that is constantly monitored and enrichment in the form of gravel pictures under all tanks, it seems like it cannot be expected that the addition of artificial plants would show such substantial improvements as seen in the study by Lee at al. 2019. Still, we would consider the average improvement of 4 % in survival rate of our enriched lines, as biologically relevant, if that could be confirmed in a larger data set.

In regards to fertility, we hypothesized that addition of artificial plants could help with stress reduction and thereby improve mating success and egg laying, but we saw no effect both with plants either added to the home-tanks or within the mating tanks. This is mostly in agreement with Woodward et al. 2019 who saw no differences in most of their settings too (Woodward, Winder, and Watt 2019). In contrast Wafer at al reported a strong increase of egg production due to structural enrichment (Wafer et al. 2016). One reason behind this could be a difference in the initial stress level. Meaning, if the stress level was already low under standard conditions, it might not be significantly reduced any further. When Sen Sarma et al. 2023 investigated the effect of stocking densities on stress levels, they saw high cortisol levels at a very low density of 1 fish/L. The cortisol levels were strongly reduced at 3 or 6 fish/L which reflects better the situation in our husbandry (Sen Sarma et al. 2023). In addition, Wafer et al. 2016 observed a different effect of grass or leaves on egg numbers in correlation with the age of fish, when comparing zebrafish aged from 110 to 180 dpf (Wafer et al. 2016). We did not see such a correlation with our fish that were aged 120 to 270 dpf but only a linear increase of average clutch size with age for all conditions tested (data not shown). In our own experience, young and healthy fish (3-12 months post fertilization, mpf) mate easily and very successful even in barren environments. But for older fish or lines with higher degrees of inbreeding that would otherwise be less fertile, enrichment in the form of gravel pictures or providing areas with shallow and deeper water in the mating tanks can improve chances for a successful mating. Therefore, it would be interesting to investigate in the future if addition of artificial plants could help with health and fertility in older fish (> 15 mpf).

So far, the strongest evidence in favor of structural enrichment comes from place preference experiments that showed that zebrafish would rather occupy higher structured areas than barren tanks (Stevens, Reed, and Hawkins 2021). At the same time this highly social fish species displays strong shoaling behavior which can only be performed in open water (Miller and Gerlai 2011). By observing wild caught zebrafish in a large arena, it has been suggested that the preference for more vegetation could even outweigh the desire for larger groups (Ghoshal and Bhat 2021). But in contrast to the vast ponds and rivers of the natural habitat, the current commercial standard of tank sizes of up to 10 L and compared to nature high holding densities can be challenging to provide enough space to allow for both. Therefore, we wanted to examine the long-term effects of structural enrichment in regards of place preference in our holding tanks. In addition, an animal facility can be a busy place and we examined how human presence might affect this preference. We found that under standard conditions without enrichment the fish had no preference for the left or right tank side but instead preferred the open water in the middle over the areas with side walls. When artificial leaves were placed at the left side of the tank, fish were significantly less likely to be present in that area but instead moved more to the middle compartment or the previously less favored right wall. This effect was seen both with and without human presence but more pronounced in the first case. Here, even a distant perceived human presence was sufficient to attract the fish closer to the front out of curiosity or in expectation of feeding. The extent of this effect depended on holding densities as well, with increasing fish numbers correlating to their presence close to plants. We concluded that only when fish seemed unwilling to further increase their shoal cohesion and needed to maintain an average distance to all other fish in their vicinity as well, they entered enriched areas. When undisturbed, the distribution of fish with enrichment was more similar to standard conditions without enrichment. A possible explanation could be that undisturbed fish swim slower on average and are therefore able to move closer to the artificial plants or were occupying more often rear parts of the tanks, resulting in a less pronounced change of place preference. The remaining difference to standard conditions might be just the physical volume the plants occupy and not an induced change in fish behavior. This needs to be investigated in more detail in the future. Still, in contrast to the initial working hypothesis, the artificial plants were clearly not able to attract more fish to their closer vicinity but in the best case fish were indifferent to their presence and in the worst case fish were displaced in areas usually avoided due to restriction of free swimming space.

Further research is needed on how the design and position of the structural enrichment can be improved to be more attractive for fish at holding densities that are typical for husbandries. These should also reduce the likelihood of hierarchical structures developing among the fish or at least provide sufficient hiding places from dominant or aggressive fish. Based on our findings of shoaling place preference, fish seemed to prefer open water over areas close to tank walls and least the plant enriched areas. Still, individual fish might profit from enrichment either during upbringing, before shoaling behavior is established or when hiding space is needed to escape aggression of their siblings. Furthermore, even when fish only occasionally interact with structural enrichment, it could still be sufficient for their cognitive improvement, which we have not tested (von Krogh et al. 2010; Gatto et al. 2022).

In summary, we think that in previous publications lack of standardization and major differences in husbandry procedures and systems resulted in different outcomes or extent of the effects of structural enrichment. We found that establishing structural enrichment in our facility was low cost and had no detrimental effects on the fish. Despite a promising trend on improved survival for rearing fish, we found no significant benefits in our readouts. Here, additional data with a larger sample size would be needed to determine these more subtle effects which our small facility could not provide yet. Therefore, we would encourage other facilities to consider new applicable forms of enrichment and report their findings to further an open discussion regarding good practices. Most importantly fish behavior in the form of aggression or place preference needs to be studied in long-term observations in more detail in the future. As previous research on older animals suggested, a more structured environment could be especially important to maintain health and cognitive functions at advanced age. Another future challenge will be to balance the preference for enrichment, open water and optimal holding density in comparatively small husbandry tanks. In the long-term improved forms of husbandry systems should be considered that allow for more space to include both.

## Supporting information

Supplemental Image 3

## Author Contributions

AK, RB, IB, AS and NO performed the experiments. NO performed statistical analysis of data. JW and NO contributed to experimental design. NO provided the first draft of the manuscript. JW added major revisions. All authors read and approved the final manuscript.

## Acknowledgements

We thank our animal care takers for maintenance of BfR zebrafish facility, providing excellent husbandry conditions and their support in collecting the data. We thank the whole team of the Aqua facility for helpful suggestions and discussions.

## Figure Legends

Supplement 1: table PubMed Search

Supplement 2: raw data tables raising fish and mating

Supplement 3: gravel picture for printing

