## Supplementary figures and images for "Zebrafish facility report on implementation of artificial plants as structural enrichment"

### Supplemental Image 3

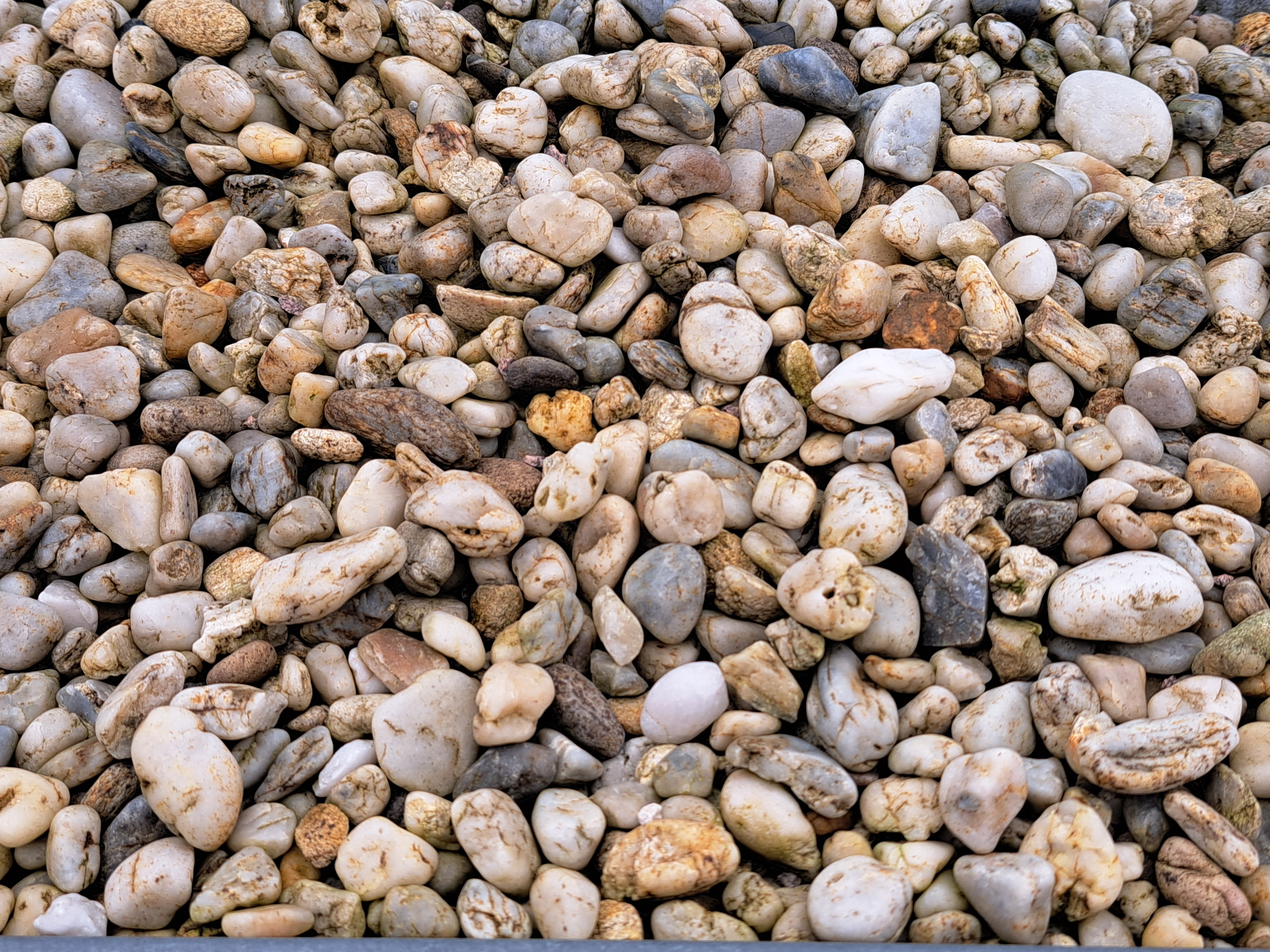
